# 6mA demethylase Mi-NMAD-1 modulates nematode development and parasitism through the transcription factor Mi-nhr-118

**DOI:** 10.64898/2025.12.16.694795

**Authors:** Dadong Dai, Yucheng Liao, Dexin Bo, Shahid Siddique, Yayi Zhou, Boyan Hu, Shurong Zhang, Yali Zhang, Pengfei Zhu, Donghai Peng, Ming Sun, Jinshui Zheng

**Affiliations:** National Key Laboratory of Agricultural Microbiology, Huazhong Agricultural University, Wuhan 430070, China; Hubei Hongshan Laboratory, Wuhan 430070, China; Department of Entomology and Nematology, University of California Davis, One Shields Avenue, Davis, CA 95616, USA

## Abstract

N6-methyladenine (6mA) DNA methylation has emerged as an important epigenetic mark in eukaryotes, but its biological role in plant-parasitic nematodes remains largely unexplored. Here, we demonstrate that 6mA methylation is essential for the embryonic development and parasitic ability of *Meloidogyne incognita*. Silencing of the 6mA demethylase genes *minmad-1* significantly reduced egg hatching, revealing that 6mA-mediated regulation is required for nematode development. Transcriptome analyses showed that *minmad-1* knockdown downregulated genes involved in embryogenesis, RNA biosynthesis, and cell differentiation, suggesting a critical role of 6mA in developmental gene regulation. Among the transcription factors affected by *minmad-1* silencing, *Mi-nhr-118*, a nuclear hormone receptor orthologous to *C. elegans nhr-118*, was identified as a key regulator of embryonic development. Functional assays confirmed that RNAi or host-induced gene silencing (HIGS) of *Mi-nhr-118* led to developmental arrest, reduced gall and egg mass formation, and impaired infectivity across multiple *Meloidogyne* species. ChIP-seq analysis revealed that *Mi-nhr-118* directly binds to promoters of genes associated with cellular regulation and developmental signaling. Several downstream targets, including transcription factors (Mi_23621.1, Mi_39647.1) and genes implicated in metabolic and signaling processes (Mi_03067.1, Mi_08398.1), were shown to be essential for both embryogenesis and parasitism. Functional enrichment analyses of *Mi-nhr-118* target genes indicated suppression of immune-related signaling, carbohydrate recognition, and glycoprotein biosynthetic pathways following gene silencing. Collectively, these findings establish *Mi-nhr-118* as a central regulator linking 6mA-mediated epigenetic control to development and parasitism in *M. incognita*.

## Introduction

Epigenetic modifications are molecular mechanisms that regulate gene expression and phenotype without altering the underlying DNA sequence ^1^. These modifications influence transcriptional activity through multiple processes, including DNA methylation, histone methylation and acetylation, chromatin remodeling, and the action of non-coding RNAs ^2,3^. Beyond the canonical 5-methylcytosine, N6-adenine methylation (6mA) has emerged as a bona fide eukaryotic DNA modification implicated in development, stress responses, and transgenerational inheritance in diverse taxa, including the nematodes ^4–7^. In *C. elegans*, 6mA has been functionally implicated in DNA modification processes ^8^, intergenerational inheritance of mitochondrial DNA mutations ^9^, and the regulation of aging ^10^. However, how 6mA is wired into transcription-factor (TF) networks to coordinate complex biological programs in obligate plant pathogens remains largely unknown.

Root-knot nematodes (RKNs) of the genus *Meloidogyne* are among the most destructive plant parasites worldwide, infecting thousands of hosts and causing major yield losses ^11–15^. Species within the *M. incognita* group (MIG) combine polyploidy with obligate parthenogenesis ^16,17^. These features limit adaptive genetic recombination and point to epigenetic plasticity as a plausible route to rapid reprogram development and parasitism ^16^. Pinpointing the molecular conduits that translate epigenetic marks into gene regulatory outputs is therefore central to understanding how these nematodes execute the coupled cascade from development to infection. In our previous work, we confirmed the presence of 6mA DNA methylation in *M. incognita* using multiple analytical approaches and identified its demethylase, Mi-NMAD-1/2, through integrated bioinformatic and experimental analyses ^5^. Our results indicate that the 6mA demethylase modulates the expression of nematode virulence factors, and host-induced gene silencing (HIGS) of *minmad-1/2* significantly suppresses the nematode’s ability to infect host plants ^5^. However, two key questions remained unanswered: (i) which downstream transcriptional regulators are directly under the control of Mi-NMAD-1–mediated demethylation, and (ii) how such regulators connect developmental programs with parasitic competence in *M. incognita*.

In this study, by comparing the transcriptomic profiles of *minmad-1* RNAi nematodes with those undergoing normal egg-to-J2 development, we identified a nuclear hormone receptor, Mi-nhr-118, as a demethylase-responsive transcription factor homologous to *C. elegans nhr-118* and conserved across *Meloidogyne* species. Mi-NMAD-1 maintains normal *Mi-nhr-118* expression by removing 6mA methylation, thereby ensuring proper embryonic development. Silencing *Mi-nhr-118* in *M. incognita* embryonic cells markedly impaired embryogenesis, whereas RNAi at the J2 stage significantly reduced the nematode infection rate in host plants. Consistently, transgenic plants expressing *Mi-nhr-118* double-stranded RNA showed increased resistance to *M. incognita*, *M. javanica*, and *M. arenaria*. By integrating RNA sequencing after *Mi-nhr-118* silencing with chromatin immunoprecipitation followed by sequencing (ChIP-seq) of Mi-nhr-118 binding sites, we further show that this transcription factor regulates a suite of downstream genes involved in embryogenesis, sugar transport, signal transduction, and parasitism. These findings provide a mechanistic link from 6mA demethylation to transcription factor activation and ultimately to nematode infectivity. In summary, our work reveals a previously unrecognized epigenetic and transcriptional regulatory pathway in a parthenogenetic plant-parasitic nematode and identifies practical molecular targets for HIGS-based control strategies.

## Result

### 6mA DNA methylation is involved in the embryonic development of *M. incognita*

Our previous study demonstrated that silencing demethylase genes significantly reduced the virulence of *M. incognita* group (MIG) during host plant infection ^5^. To further explore the molecular mechanisms by which the 6mA demethylase regulates the virulence of *M. incognita*, we performed RNA-seq on eggs collected from the *minmad-1/2*- and *Mi_23849.1*-HIGS low-gall group, as well as from the ds*gfp* control group. Afterwards, a total of 1,122, 767, and 407 differentially expressed genes (DEGs) were identified from eggs of *M. incognita* infecting *minmad-1/2*-, and *Mi_23849.1*-HIGS transgenic plants (Supplementary Fig. 1), respectively. The Pfam enrichment analysis of DEGs showed that the methyltransferase was enriched and down-regulated in *minmad-1/2* knockdown groups (Fig. 1A, B), but not in *Mi_23849.1* knockdown group (Supplementary Fig. 2). Furthermore, the significantly down-regulated DEGs in the demethylase RNAi group were enriched in multiple biological pathways such as GPCR receptor family, sugar transporter MFS-5, lipolysis regulator Seipin, and Collagen (Fig. 1A, B). These findings indicate that the demethylases modulate the parasitic capacity of *M. incognita* via multiple molecular pathways.

**Figure 1.**
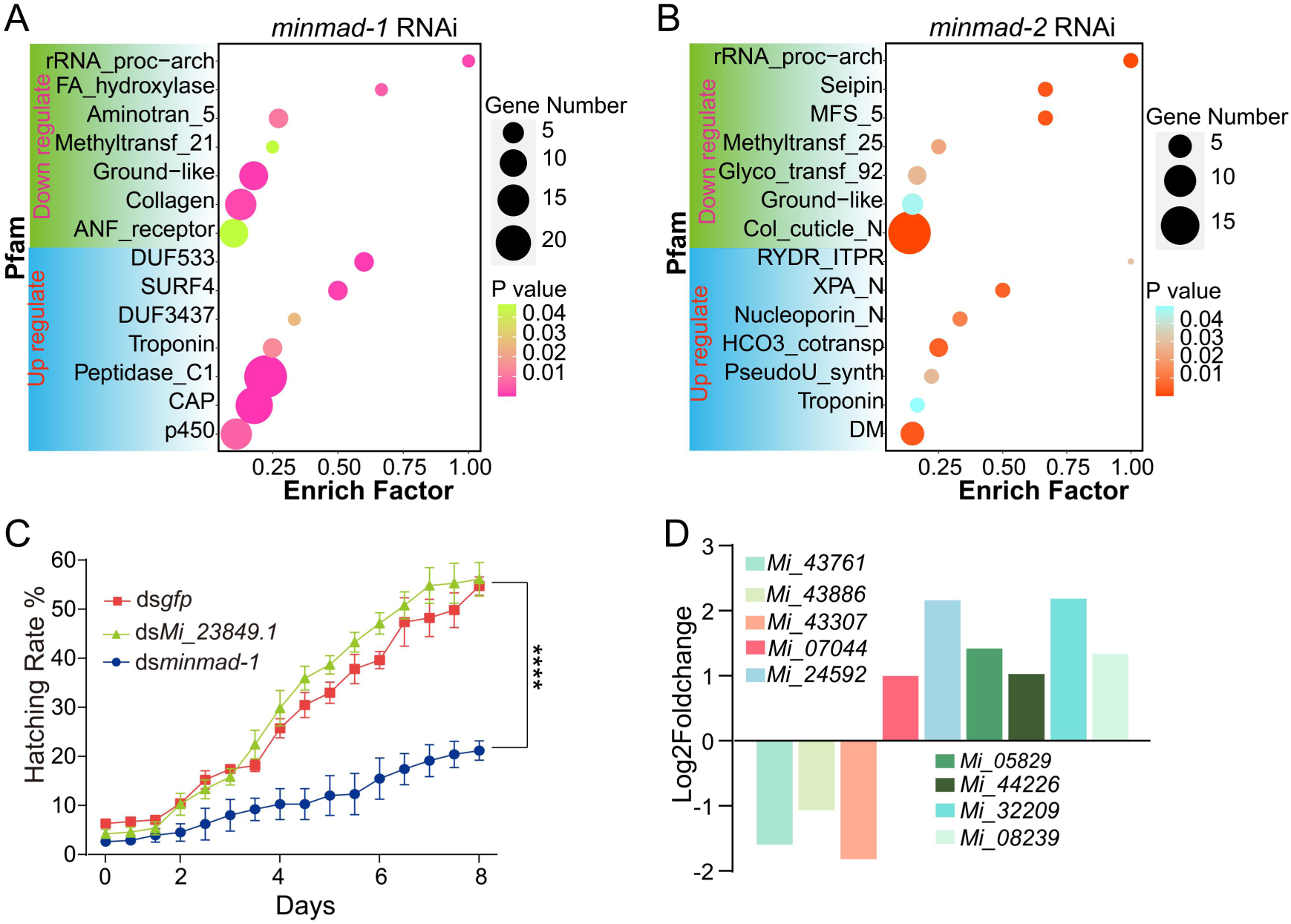
*minmad-1* RNAi inhibited embryonic development in *M. incognita*. (A) Pfam enrichment analysis of DEGs in *M. incognita* eggs from *minmad-1*-HIGS lines revealed significant down-regulation of genes associated with methyltransferases and collagen. (B) Pfam enrichment analysis of DEGs in *M. incognita* eggs from *minmad-2*-HIGS lines revealed significant down-regulation of genes associated with methyltransferases and sugar transporter MFS-5. (C) In vitro RNAi targeting *minmad-1* in *M. incognita* embryonic cells caused developmental arrest, significantly reducing the proportion of embryos that developed into J2s. (D) Nine differentially expressed transcription factors (TFs) overlapped between the *minmad-1* RNAi and egg-to-J2 datasets, among which three were down-regulated in the *minmad-1* RNAi treatment but up-regulated during normal egg-to-J2 development.

To comprehensively investigate the molecular role of 6mA DNA methylation in *M. incognita*, particularly in developmental processes, we silenced the demethylase *minmad-1* in embryo and examined the resulting phenotypes (Supplementary Fig. 3). Strikingly, RNAi of *minmad-1* led to a significant reduction in hatching rate (Fig. 1C), indicating that 6mA is essential for embryonic development. To uncover the underlying molecular mechanisms, we performed RNA-seq on embryos after *minmad-1* knockdown. A total of 1,916 DEGs were identified when comparing *minmad-1* RNAi with the ds*gfp* treatment control. Gene Ontology (GO) enrichment analysis revealed that these DEGs were predominantly associated with terms related to embryogenesis, including organismal development, RNA biosynthesis, pharyngeal gland morphogenesis, apoptotic signaling, and autophagosome formation (Supplementary Fig. 4). Moreover, Pfam domain enrichment highlighted categories such as apoptosis-associated TTR-52, stem cell differentiation-related Laminin_N ^18,19^, RNA-binding protein La, extracellular receptor SRCR, and cadherin domains involved in cell-cell adhesion during development (Supplementary Fig. 4). Together, these findings demonstrate that 6mA DNA methylation plays a critical role in regulating embryonic development in *M. incognita*.

Since transcription factors (TFs) are key regulators of gene expression, we examined the DEGs described above for TFs and identified 40 candidates that were significantly altered following *minmad-1* knowdown. Moreover, since *minmad-1* knockdown markedly impaired egg hatching, thereby reducing the transition from eggs to second-stage juvenile (J2), we further intersected these 40 TFs with those DEGs during the egg-to-J2 developmental transition. This comparison yielded nine overlapping TFs. Strikingly, their expression patterns were reversed: TFs upregulated during the egg-to-J2 transition were consistently downregulated in *minmad-1* RNAi-treated eggs, in line with our expectations (Fig. 1D). We speculate that the three TFs *Mi_43761.1*, *Mi_43886.1*, and *Mi_43307.1*, which were down-regulated following *minmad-1* RNAi while up-regulated during egg-to-J2 transition, may participate in regulating the embryonic developmental process of *M. incognita*.

### Identification of *Mi-nhr-118* as a growth-related TF

To verify whether these three candidate TFs are involved in the embryonic development of *M. incognita*, we performed in vitro RNAi on embryo. The results showed that, compared with the ds*gfp* and wild-type controls, the hatching rate of embryos in the ds*Mi_43307.1* treatment group was significantly reduced, whereas ds*Mi_43761.1* and ds*Mi_43886.1* showed no significant differences (Fig. 2A, Supplementary Fig. 5). To confirm the effect of Mi_43307.1 knockdown on embryonic development, we monitored developmental progression across multiple time points and observed that the developmental arrest persisted until cell lysis occurred (Fig. 2B). Moreover, the phenomenon of developmental arrest after *minmad-1* RNAi was likewise observed in both *M. arenaria* and *M. javanica* (Supplementary Fig. 6). These findings suggest that Mi_43307.1 may participate in the demethylase-mediated regulation of embryonic development in *M. incognita*. To further investigate the relationship between the demethylase and Mi-43307.1, we examined previously generated Mi-NMAD-1 ChIP-seq data [10] and found that the demethylase directly binds to the transcription factor Mi_43307.1 at the J2 stage (Fig. 2C). We also compared the 6mA-DIP-seq profiles before and after *minmad-1* RNAi at the egg stage and found that 6mA signals became detectable on *Mi_43307.1* following *minmad-1* silencing (Supplementary Fig. 7). These data indicate that 6mA and its demethylase Mi-NMAD-1 indeed act directly on the transcription factor Mi_43307.1.

**Figure 2.**
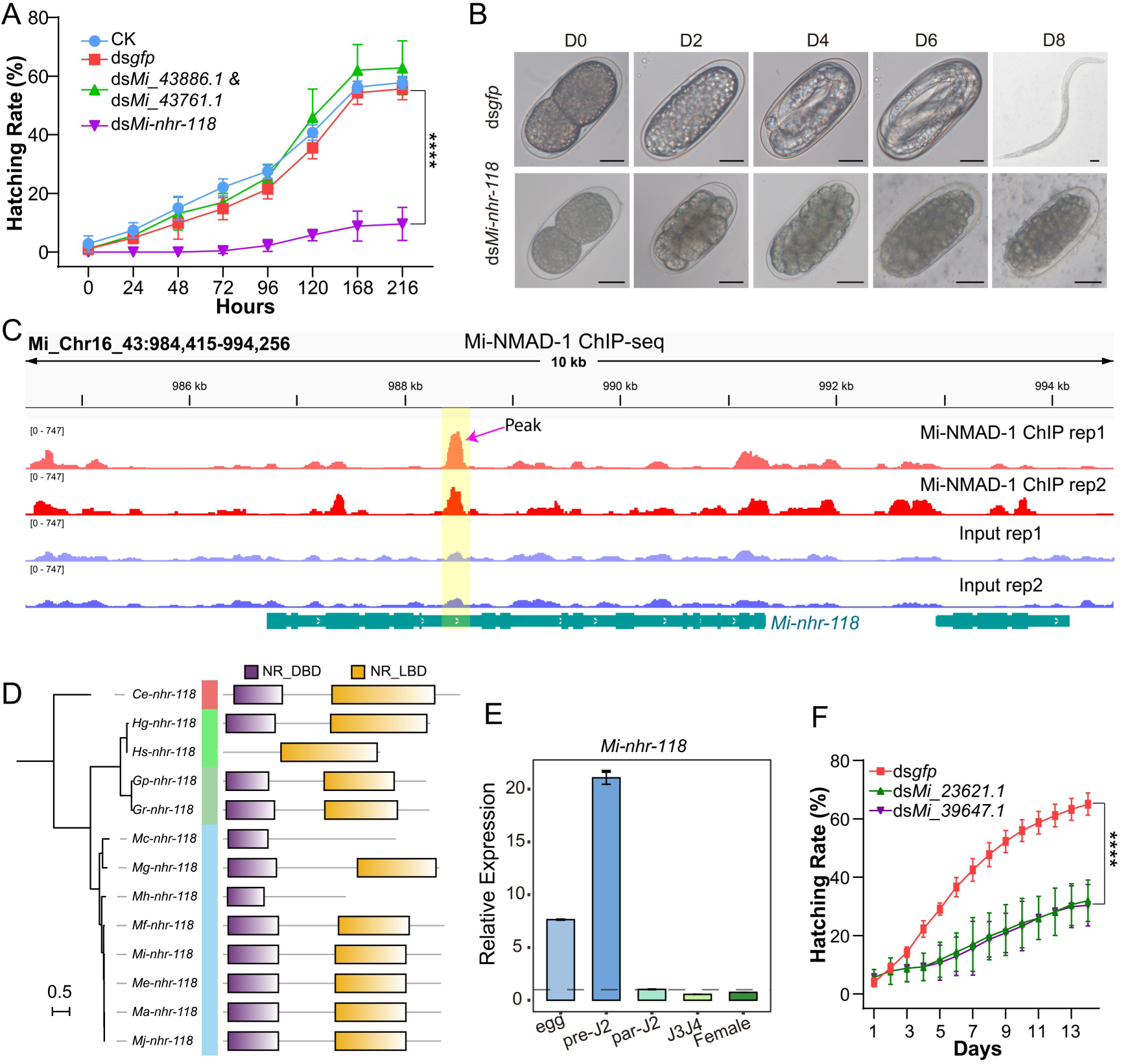
*minmad-1* regulates the expression of downstream embryogenesis-related genes through the nuclear hormone receptor transcription factor Mi-nhr-118. (A) In vitro RNAi results showed that, compared with the CK and ds*gfp* controls, silencing of *Mi-nhr-118* significantly reduced the proportion of *M. incognita* embryos developing into J2s, whereas no significant changes were observed for *Mi_43886.1* and *Mi_43761.1*. Given the high degree of sequence conservation shared by these two genes, RNAi treatment was designed to target them simultaneously. (B) RNAi targeting *Mi-nhr-118* caused developmental arrest in embryos, with a large proportion failing to develop into juveniles, whereas embryos in the ds*gfp*-treated control group developed normally. (C) ChIP-seq data of Mi-NMAD-1 at the J2 stage of *M. incognita* showed that it directly targets the nuclear hormone receptor gene Mi-nhr-118. (D) Phylogenetic analysis of Mi-nhr-118 homologs in nematodes and the conserved domains of the corresponding proteins. (E) Relative expression levels of *Mi-nhr-118* at different developmental stages. (F) In vitro RNAi of the differentially expressed transcription factors identified after *Mi-nhr-118* RNAi similarly resulted in embryonic developmental arrest in *M. incognita*.

A domain-based BLAST search against the Conserved Domain Database (CDD) revealed that Mi_43307.1 contains a DNA-binding domain (8–85 aa; accession: cl02596) and a ligand-binding domain of nuclear hormone receptors (172–280 aa; accession: cl11397) (Supplementary Fig. 8A, B). BLAST analysis against the WormBase nonredundant protein database showed that Mi_43307.1 shares high sequence similarity with *Ce-nhr-118* from *C. elegans* and was therefore designated *Mi-nhr-118*. Homologs of this gene were also widely identified in *Heterodera*, *Globodera*, and *Meloidogyne*. A phylogenetic tree constructed from protein sequences revealed that *Mi-nhr-118* clusters most closely with other *Meloidogyne* homologs (52–99% identity), particularly *Ma-nhr-118* (*M. arenaria*, 98.2%), *Me-nhr-118* (*M. enterolobii*, 98.5%), and *Mj-nhr-118* (*M. javanica*, 98.2%) (Fig. 2D). Functional studies of NHR transcription factors in *C. elegans* have shown that they play important roles in promoting cell differentiation and other developmental processes ^20^, supporting the role of *Mi-nhr-118* in embryonic development of *M. incognita*.

To further characterize *Mi-nhr-118*, we examined its transcriptional expression pattern across the developmental stages of *M. incognita* using reverse transcription quantitative PCR (RT-qPCR). The expression level at the female stage was set as the reference (value = 1) for comparison across other stages. *Mi-nhr-118* expression was significantly upregulated in eggs, pre-parasitic J2s, and post-parasitic J2s (3 days post-infection, dpi), peaking in pre-J2s with a ∼25-fold increase relative to females (Fig. 2E). In contrast, the lowest transcript levels were detected in mixed J3/J4 (12–18 dpi) and female stages (Fig. 2E). These results suggest that *Mi-nhr-118* is predominantly active during the early developmental and infective stages of *M. incognita*, implicating it in processes associated with embryogenesis and host invasion.

Then, we performed transcriptome sequencing on *M. incognita* embryos treated separately with *Mi-nhr-118* and *gfp* dsRNA for 48 hours to further elucidate the molecular function of *Mi-nhr-118*. Embryos from these two treatments were collected to construct four RNA-seq libraries. A total of 1,532 DEGs were identified, including 791 upregulated and 739 downregulated genes in *Mi-nhr-118*-RNAi embryos compared with the ds*gfp* controls (Supplementary Fig. 9A). GO enrichment analysis showed that the DEGs were significantly enriched in terms related to developmental processes, with 149 genes upregulated and 209 downregulated (Supplementary Fig. 9B). We examined the genes that were downregulated following *Mi-nhr-118* RNAi and identified 40 transcription factors among them. Further in vitro RNAi of two of these transcription factors (Mi_23621.1, Mi_39647.1) similarly resulted in arrested embryonic development and a reduced number of hatched eggs (Fig. 2F). Collectively, these results demonstrate that silencing the nuclear hormone receptor *Mi-nhr-118* disrupts the expression of multiple development-associated genes, thereby impairing embryonic development in *M. incognita*.

### *Mi-nhr-118* is essential for the embryonic development and parasitism of *M. incognita* group

Since *Mi-nhr-118* exhibited its highest expression during the J2 stage, the only infective stage, we hypothesized that this gene might also be involved in regulating the parasitic ability of *M. incognita*. After performing in vitro RNAi on J2, we observed a significant reduction in their ability to infect tomato plants, with both the number of galls and egg masses markedly lower than those in the ds*gfp* and blank control groups (Fig. 3A, B). To investigate the role of the nuclear hormone receptor *Mi-nhr-118* in the development of other life stages of nematodes, we engineered nuclear transgenic tobacco plants through host-induced gene silencing (HIGS) designed to express *dsMi-nhr-118* (Figure S10). We selected 3 transgenic plant lines expressing *Mi-nhr-118* dsRNA, along with the wild-type (WT) and ds*gfp* expression plants served as controls. Each plant was inoculated with 1,000 J2s, and the extent of parasitism was assessed by counting the number of galls and egg masses at 70 dpi. The results showed that, compared with nematodes infecting WT and ds*gfp* plants, infection of *Mi-nhr-118*-HIGS plants resulted in smaller galls, and no multiple generations of nematodes developed within the galls (Fig. 3C). Subsequently, we quantified the average numbers of galls and egg masses per plant. Compared with the control group, the *Mi-nhr-118*-HIGS plants showed reductions of 36.39–55.67% in gall numbers and 30.78–44.57% in egg masses (Fig. 3D, E). In addition to producing fewer galls, the transgenic plants also remained visibly healthier after nematode infection, displaying greater plant height than the controls (Supplementary Fig. 11). We also evaluated the hatching rate of eggs laid by nematodes that had fed on *Mi-nhr*-118-HIGS plants. Compared with the control group, their hatching rate decreased by 65.48% (Supplementary Fig. 12), which is consistent with the results obtained from the in vitro embryonic RNAi assays. Since the nucleotide sequence of *Mi-nhr-118* is highly conserved among MIG species, we next tested whether this transcription factor also plays an important role in *M. arenaria* and *M. javanica*. Using the same *Mi-nhr-118*-HIGS transgenic lines, we assessed the infectivity of *M. arenaria* and *M. javanica*. The results showed that the *Mi-nhr-118*-HIGS plants also exhibited strong resistance to *M. arenaria* and *M. javanica*, with both galls and egg mass numbers significantly reduced compared with the controls (Fig. 3F, G; Supplementary Fig. 13). These results indicate that *Mi-nhr-118* plays a critical role in regulating both nematode development and parasitism.

**Figure 3.**
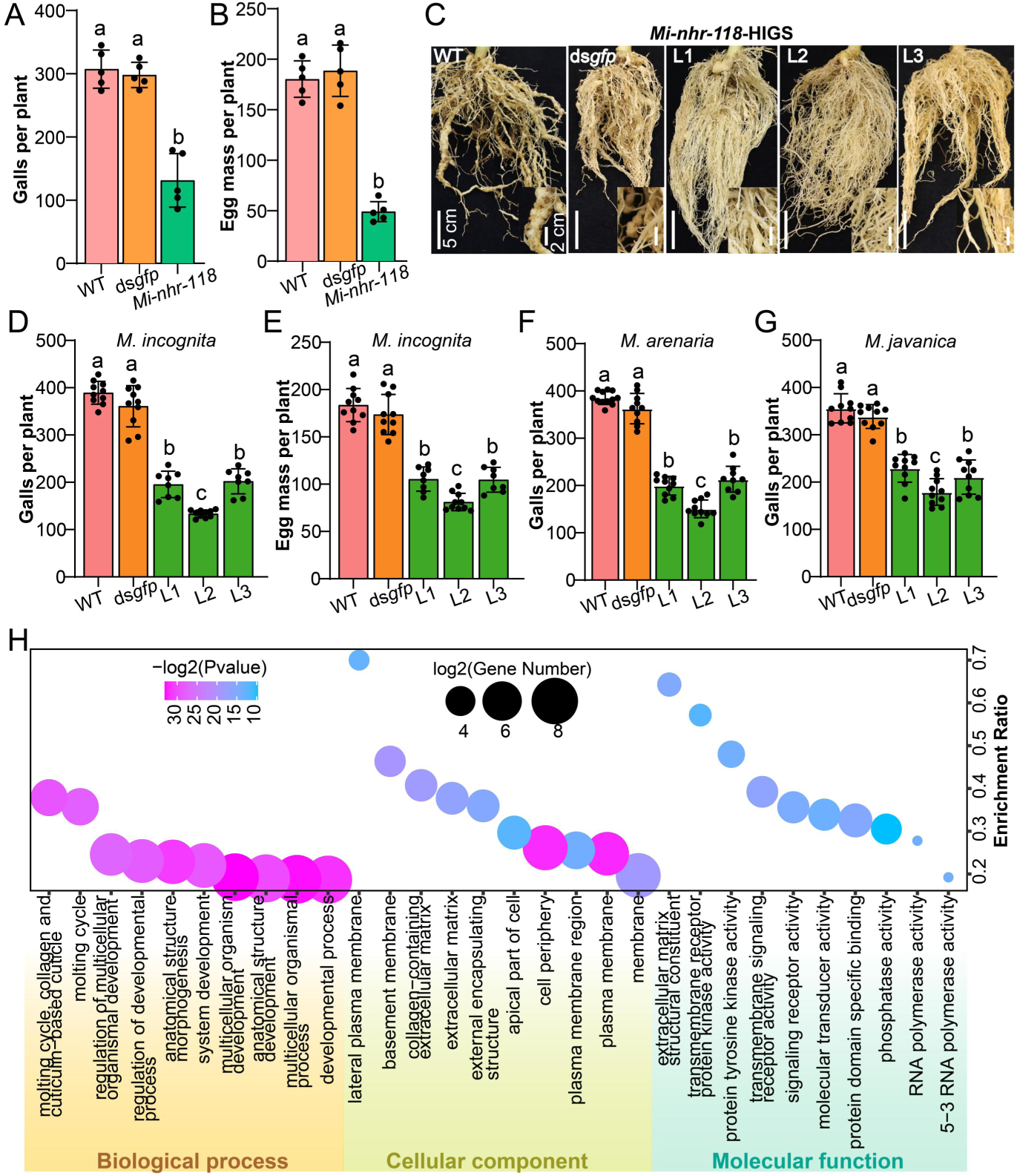
*Mi-nhr-118*-HIGS significantly enhanced the resistance of transgenic plants to *M. incognita group* (MIG) nematodes. (A) In vitro RNAi targeting *Mi-nhr-118* in *M. incognita* J2s significantly reduced the number of galls formed on host plant roots. (B) In vitro RNAi targeting *Mi-nhr-118* in *M. incognita* J2s significantly reduced the number of egg mass on host plant roots. (C) *Mi-nhr-118*-HIGS significantly reduced the size of galls induced by *M. incognita* infection. In contrast to the large, multi-generation galls observed in the CK and ds*gfp* groups, most galls formed on *Mi-nhr-118*-HIGS plants were small and contained only single infections. (D and E) *Mi-nhr-118*-HIGS significantly reduced the number of galls and egg mass formed by *M. incognita* infection in transgenic plants. L1, L2, and L3 represent the three transgenic lines with the highest expression levels that we selected. (F) *Mi-nhr-118*-HIGS significantly reduced the number of galls formed by *M. arenaria* infection in transgenic plants. (G) *Mi-nhr-118*-HIGS significantly reduced the number of galls formed by *M. javanica* infection in transgenic plants. Since the large galls in the control group could not be accurately counted, the actual number of galls was likely much higher than the recorded value. (H) GO functional enrichment analysis of down-regulation DEGs from J3/J4 stages of *Mi-nhr-118*-HIGS nematodes revealed numerous genes involved in development, molting, and RNA polymerase activity. Lowercase letters denote significant differences (*p* < 0.05, one-way ANOVA with Tukey’s test).

To further investigate the molecular basis underlying the reduced parasitic ability of nematodes after *Mi-nhr-118* RNAi, we performed RNA-seq on the J3/J4-stage nematodes collected from *Mi-nhr-118*-HIGS transgenic plants. A total of 9,179 DEGs were identified between the *Mi-nhr-118*-HIGS and ds*gfp* control groups, including 5,117 upregulated and 4,062 downregulated genes. GO enrichment analysis of these DEGs revealed that the down-regulated genes were primarily associated with multicellular organismal development, anatomical structure morphogenesis, biological regulation, and molting-related processes (Fig. 3H). Surprisingly, the upregulated DEGs were mainly enriched in reproductive processes such as chromosome segregation, gamete generation, sperm motility, and male meiotic nuclear division (Supplementary Fig. 14), suggesting that *Mi-nhr-118* may also participate in the molecular pathways regulating sex differentiation in *M. incognita*. These findings suggest that *Mi-nhr-118* is essential for maintaining proper developmental regulation and cuticle remodeling required for nematode growth and parasitic adaptation, and that its silencing disrupts these processes while aberrantly activating reproductive pathways, thereby reducing the infectivity of *M. incognita*.

### Identification of *Mi-nhr-118* binding sites and regulatory targets

To further investigate the direct target genes regulated by *Mi-nhr-118* and gain deeper insights into its functional characteristics, we generated a *Mi-nhr-118* antibody. Western blot results showed that the *Mi-nhr-118* antibody specifically recognized the target protein in both the egg and J2 stages (Supplementary Fig. 15A). We constructed two biological replicates of ChIP-seq libraries for both the egg and pre-J2 stages of *M. incognita*. We identified 1,780 peaks in the egg stage and 6,982 peaks in the J2 stage. Notably, *Mi-nhr-118* peaks were predominantly enriched in promoter regions, accounting for 75.01% (1,336/1,780) in eggs and 51.49% (3,596/6,982) in J2s (Figure 4A). From these peaks, a total of 1,687 and 5,527 genes were annotated, comprising 417 genes encoding secreted proteins and 305 encoding transcription factors.

**Figure 4.**
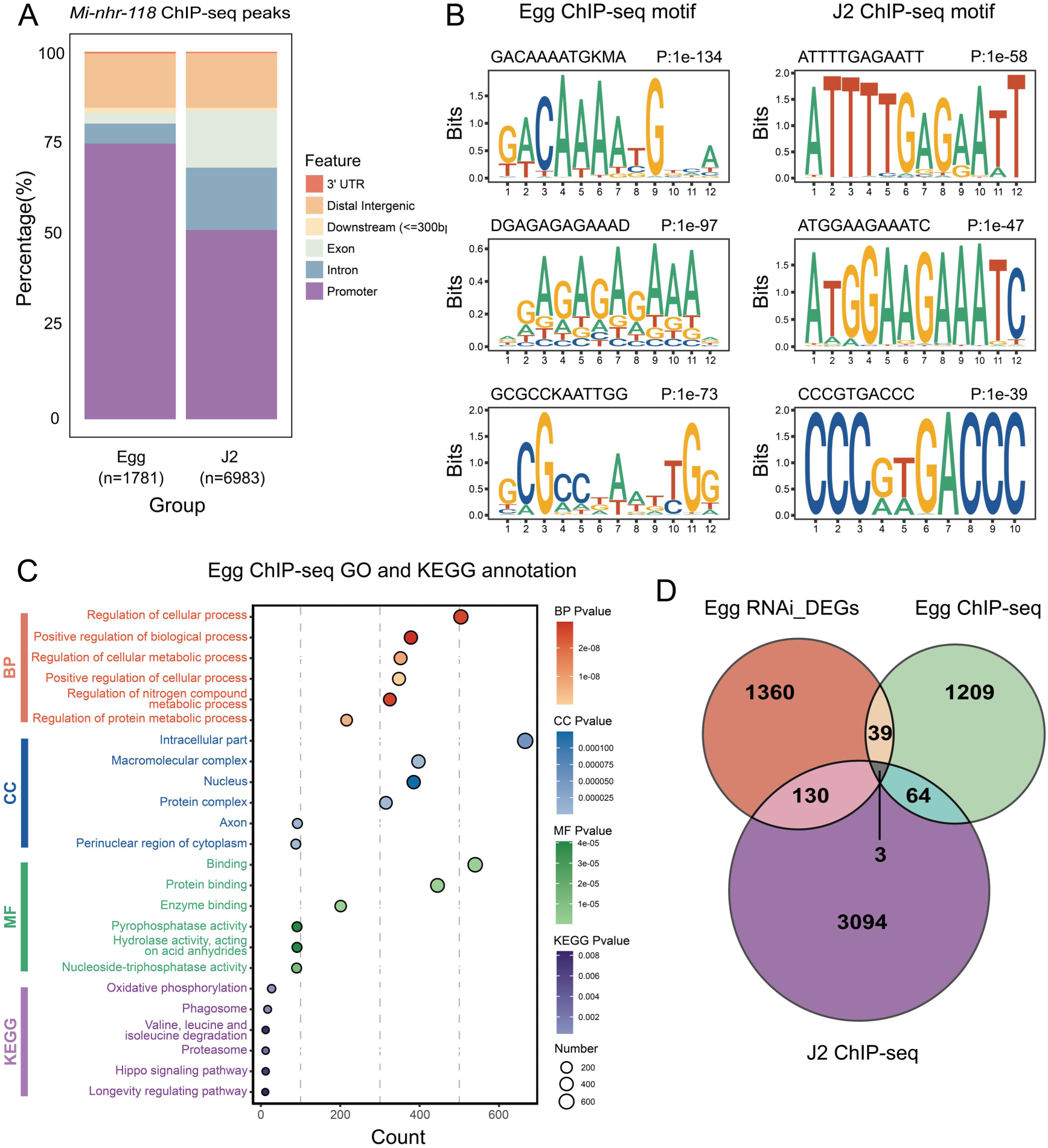
ChIP-seq analysis using a Mi-nhr-118-specific antibody was performed to identify its direct target genes. (A) Annotation of the genomic distribution characteristics of peaks obtained from Mi-nhr-118 ChIP-seq analysis. (B) Motif enrichment analysis of peaks obtained from Mi-nhr-118 ChIP-seq revealed a significant enrichment of NHR-type motifs. (C) GO and KEGG functional enrichment analyses were performed on genes annotated from Mi-nhr-118 ChIP-seq data at the egg stage. (D) Overlap analysis was performed between genes annotated from Mi-nhr-118 ChIP-seq at the egg and J2 stages and the DEGs identified after *Mi-nhr-118* RNAi in embryos.

We identified the consensus binding motif of *Mi-nhr-11*8 from the ChIP-seq dataset by using the *de novo* motif discovery algorithm in HOMER. Notably, HOMER motif enrichment analysis revealed that the *nhr-6* motif was the most significantly enriched among peaks in the egg group (Figure 4B). The *nhr-6* gene is known to be essential for regulating both cell proliferation and differentiation during nematode organ development ^21^. GO and KEGG enrichment analyses of *Mi-nhr-118* ChIP-seq peaks from eggs showed that target genes were primarily associated with the regulation of cellular and metabolic processes (Figure 4C). In contrast, *Mi-nhr-118* ChIP-seq data from J2s were mainly enriched in pathways related to microtubule polymerization and depolymerization (Supplementary Fig. 15B). We performed a comparative analysis between the list of *Mi-nhr-118* target genes identified across different developmental stages and the DEGs identifed by *Mi-nhr-118* RNAi in embryo based on RNA-seq data (Figure 4D). This analysis revealed 42 genes that were both directly bound by the nuclear hormone receptor *Mi-nhr-118* in vivo and transcriptionally regulated during embryogenesis. In addition, 67 Mi-nhr-118 binding genes were shared between the egg and J2 stages, suggesting that *Mi-nhr-118* controls a subset of common target genes across these critical developmental transitions.

### Functional characterization of *Mi-nhr-118* target genes involved in the development and parasitism of *M. incognita*

To assess the role of *Mi-nhr-118* target genes in the development of *M. incognita*, we performed in vitro RNAi assays to validate their functions. Embryos dissected from female nematodes were individually exposed to double-stranded RNA (dsRNA) corresponding to each of the 42 candidate genes. The results showed that RNAi of 20 genes led to a significant reduction in egg hatching rate (Fig. 5A; Supplementary Fig. 16). Among them, silencing of *Mi_03067.1* and *Mi_08398.1* caused particularly strong inhibition of embryonic development, resulting in a 45–48% decrease in egg hatching rate compared with the control group (Fig. 5A). Homology-based BLAST analysis further identified orthologs of *Mi_03067.1* and *Mi_08398.1* in *C. elegans* (Supplementary Fig. 17). Both *H17B01.1* (ortholog of *Mi_03067.1*, predicted to enable hexose transmembrane transporter activity) and *W01A8.8* (ortholog of *Mi_08398.1*, predicted to enable hormone activity) were found to be expressed during the embryonic stage of *C. elegans*.

**Figure 5.**
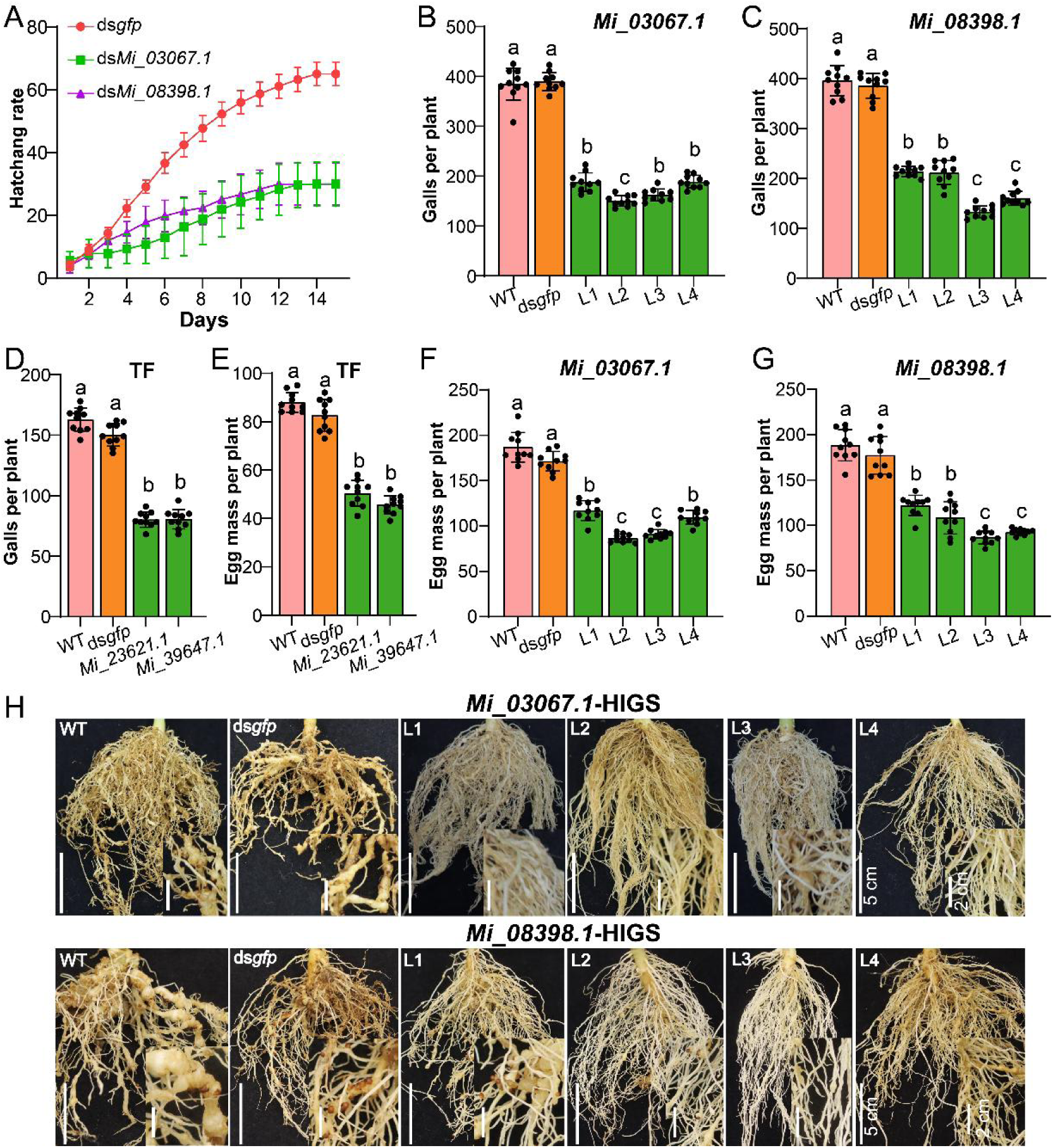
The target genes of Mi-nhr-118 play important roles in the growth and development of *M. incognita*. (A) Mi-nhr-118 target genes associated with hexose transmembrane transporter and hormone activities were also found to cause embryonic developmental arrest upon RNAi-mediated silencing. (B-D) Transgenic HIGS plants targeting the *Mi-nhr-118* downstream genes *Mi_03067.1*, *Mi_08398.1*, TF *Mi_23621.1*, and TF *Mi_39647.1* exhibited a significant reduction in the number of galls formed after *M. incognita* infection compared with the control plants. (E-G) Transgenic HIGS plants targeting the *Mi-nhr-118* downstream genes *Mi_03067.1*, *Mi_08398.1*, TF *Mi_23621.1*, and TF *Mi_39647.1* exhibited a significant reduction in the number of egg mass formed after *M. incognita* infection compared with the control plants. Since the large galls in the control group could not be accurately counted, the actual number of galls was likely much higher than the recorded value. (H) In contrast to the control plants, the *Mi-nhr-118* target gene HIGS lines failed to produce large root galls resulting from multiple infection cycles at 70 dpi. Lowercase letters denote significant differences (*p* < 0.05, one-way ANOVA with Tukey’s test).

Finally, we investigated whether the *Mi-nhr-118* downstream genes, including the two transcription factors identified above, also contribute to nematode parasitism. We examined the expression patterns of these genes across different developmental stages and found that, in addition to the embryonic stage, they were also highly expressed during the infection stage (Supplementary Fig. 18). To this end, we generated transgenic tobacco plants expressing dsRNA constructs targeting the downstream genes *Mi_03067.1* and *Mi_08398.1*, as well as the transcription factors *Mi_23621.1* and *Mi_39647.1*. These plants were inoculated with 500 or 1,000 J2s, and infection parameters were evaluated at 70 dpi. Compared with wild-type and ds*gfp* control plants, host-induced gene silencing markedly reduced *M. incognita* infection, as indicated by a significant decrease in gall formation (Figure 5B–D). Moreover, the average number of egg masses per plant was reduced by 33.9–47.4% (Figure 5F–G). Furthermore, compared with the control plants, which developed large galls after 70 dpi as a result of multiple infection cycles, most target gene HIGS plants formed only single, small galls (Fig. 5H), indicating that these transgenic lines effectively inhibited nematode growth, development, and reproduction following infection establishment. To gain a more comprehensive understanding of the molecular functions through which *Mi-nhr-118* regulates nematode parasitism, we performed RNA-seq analysis on females and eggs collected from *Mi_23621.1*-HIGS and *Mi_39647.1*-HIGS plants after nematode infection. The analysis identified 548 and 780 DEGs in eggs, and 7,358 and 3,430 DEGs in females, respectively. We performed functional enrichment analysis on the DEGs that were downregulated in females of *Mi_23621.1*-HIGS, and the results revealed a significant overrepresentation of terms related to immune activation and cell migration (Supplementary Fig. 19), suggests that the TFs Mi_23621.1 may play an important role in response to host immune system in nematodes. Moreover, GO terms associated with carbohydrate recognition and signaling activation, including carbohydrate binding, galactose binding, oligosaccharide binding, and protein kinase activator activity, were also significantly enriched (Supplementary Fig. 19), indicating a strong capacity for sugar recognition and signal transduction. Besides, downregulated DEGs of *Mi_39647.1*-HIGS revealed significant enrichment of glycoprotein and proteoglycan biosynthetic processes, including dolichol- and polyprenol-mediated oligosaccharide assembly and isoprenoid metabolism (Supplementary Fig. 19), suggesting a suppression of glycosylation and membrane-associated biosynthetic activities that may impair cellular communication and host interaction in *M. incognita*. These results suggest that the *Mi-nhr-118* target genes play crucial roles not only in nematode development but also in establishing and maintaining successful parasitism within the host.

## Discussion

Polyploid RKNs, exemplified by *M. incognita*, represent a group of parthenogenetic agricultural pathogens that pose serious threats to global crop productivity ^22^. Despite their profound agronomic impact, fundamental biological research on these nematodes remains comparatively limited. Owing to their parthenogenetic mode of reproduction, the absence of genetic recombination and exchange typical of sexual species suggests that epigenetic mechanisms may play pivotal roles in regulating their life cycle ^23^. In our previous work, we demonstrated the presence of DNA 6mA methylation in *M. incognita* and identified Mi-NMAD-1/2 as its 6mA demethylase ^5^. Silencing Mi-NMAD-1 expression during the embryo stage markedly impaired embryonic development and subsequent life-stage transitions, indicating that methylation-based epigenetic regulation is essential for developmental control in *Meloidogyne* spp. Exploring the relationship between DNA methylation and the regulation of pathogen development and virulence may provide new insights and resources for the development of sustainable, environmentally friendly strategies for pathogen control in the future.

In recent years, many studies have uncovered numerous relationships between epigenetics and pathogen virulence. In the wheat pathogen *Zymoseptoria tritici*, effector gene clusters are enriched in genomic regions marked by H3K9me3 or H3K27me3, and multi-omics analyses have revealed that these histone marks are dynamically remodeled during infection in parallel with changes in gene expression ^24^. Deletion of the H3K27 methyltransferase KMT6 in *Fusarium graminearum* results in growth defects and loss of fertility, accompanied by marked alterations in the expression of virulence-associated metabolites and pathogenicity-related genes ^25^. In *Botrytis cinerea*, the DNA 6mA methyltransferase BcMETTL4 influences genome-wide 6mA levels and fungal virulence, and its functional mutation leads to a reduction in pathogenicity ^26^. In *Phytophthora* species, small RNA–mediated epigenetic silencing can act over long distances and heritably suppress effector gene expression ^27^. In addition, extensive studies have shown that epigenetic regulation also plays vital roles in organismal development and other biological processes. For example, in *C. elegans*, epigenetic mechanisms have been shown to regulate both development and aging ^28^. Histone modifications in RKNs have also been reported to be associated with developmental regulation ^29^. However, our understanding of the molecular mechanisms underlying epigenetic regulation of development and virulence in pathogens remains limited, particularly regarding how these mechanisms modulate complex downstream gene networks. As key regulators of gene expression, transcription factors are likely to play important roles in these processes.

Our results show that 6mA demethylase *Mi-nmad-1* in *M. incognita* not only affects the expression of nematode virulence genes but also significantly inhibits embryonic cell development when silenced by RNAi during the embryonic stage. Further investigation revealed that the transcription factor *Mi-nhr-118* plays a key role in this regulatory process. ChIP-seq and transcriptome analyses collectively revealed that *Mi-nhr-118* directly regulates a suite of downstream genes implicated in diverse biological processes which are essential for nematode growth and parasitism. HIGS experiments further validated the importance of *Mi-nhr-118* and its target genes in parasitism, as transgenic plants expressing dsRNA constructs targeting these genes exhibited strong resistance to *M. incognita* and related species. Importantly, this study provides the mechanistic insights into how epigenetic modifications can influence transcription factor–mediated regulatory networks in PPNs. The demethylase-dependent activation of *Mi-nhr-118* parallels similar mechanisms described in other systems, where DNA or histone demethylation modulates transcription factor accessibility and binding. In summary, by generating multiple HIGS transgenic plants targeting five key genes and integrating 38 RNA-seq and 8 ChIP-seq datasets, we provide a comprehensive and multi-layered understanding of how *minmad-1*–mediated transcriptional regulation governs nematode development and parasitism.

Nuclear hormone receptor (NHR) transcription factors represent a conserved family within the phylum Nematoda, functioning as key regulators of developmental and physiological processes ^30^. In *C. elegans*, NHRs mediate diverse biological functions, including density-dependent developmental acceleration via NHR-8 and DAF-12, DNA damage–induced apoptosis through NHR-14 and CEP-1/p53, and cell differentiation via NHR-25–mediated repression of GATA factors ^20,31,32^. RNAi has been a powerful tool for studying nematode gene function, and recent work revealed that perturbation of 103 *C. elegans* NHRs uncovered a modular network in which NHR pairs co-regulate shared target genes ^33^. Beyond *C. elegans*, NHRs play critical roles in other nematodes: NHR-17 and NHR-105 are indispensable for development and viability in *Haemonchus contortus* ^34^, Dd-NHR-1 governs embryogenesis in *Ditylenchus destructor* ^35^, and Mi-NHR-48 modulates both development and infectivity in *M. incognita* ^36^. However, the mechanistic relationship between 6mA demethylation and NHR-mediated transcriptional regulation has remained unclear. In this study, we reveal that the 6mA demethylase Mi-NMAD-1 regulates embryonic development and parasitism in *M. incognita* by modulating the expression of the nuclear hormone receptor *Mi-nhr-118*. Given the parthenogenetic and obligate parasitic lifestyle of *M. incognita*, which faces strong selective pressures from diverse hosts and soil environments, the evolution of an integrated DNA methylation–NHR regulatory axis likely provides a flexible epigenetic framework for developmental control and host adaptation.

Based on these findings, we propose a mechanistic model in which the demethylase Mi-NMAD-1 removes 6mA methylation from the *Mi-nhr-118* locus in *M. incognita*, thereby enabling its proper transcription and translation into the functional transcription factor Mi-NHR-118. Once activated, Mi-NHR-118 induces the expression of its downstream target genes, thereby orchestrating not only early embryonic development but also subsequent parasitic interactions with host plants in *M. incognita* (Figure 6). Overall, this study systematically characterizes the biological functions of nuclear hormone receptors (NHRs) in RKNs and elucidates their coordinated regulatory interactions with epigenetic modifications. We demonstrate that NHRs govern embryonic development and play pivotal roles in regulating nematode parasitism and adaptation to host plants. These findings highlight the potential of simultaneously targeting NMAD-1 and NHRs to disrupt nematode development and infection processes, providing a conceptual framework for the design of target-specific ovicidal or anti-nematode compounds. Such strategies identify promising molecular targets for breeding nematode-resistant crops and ultimately contribute to sustainable agriculture and global food security.

**Figure 6.**
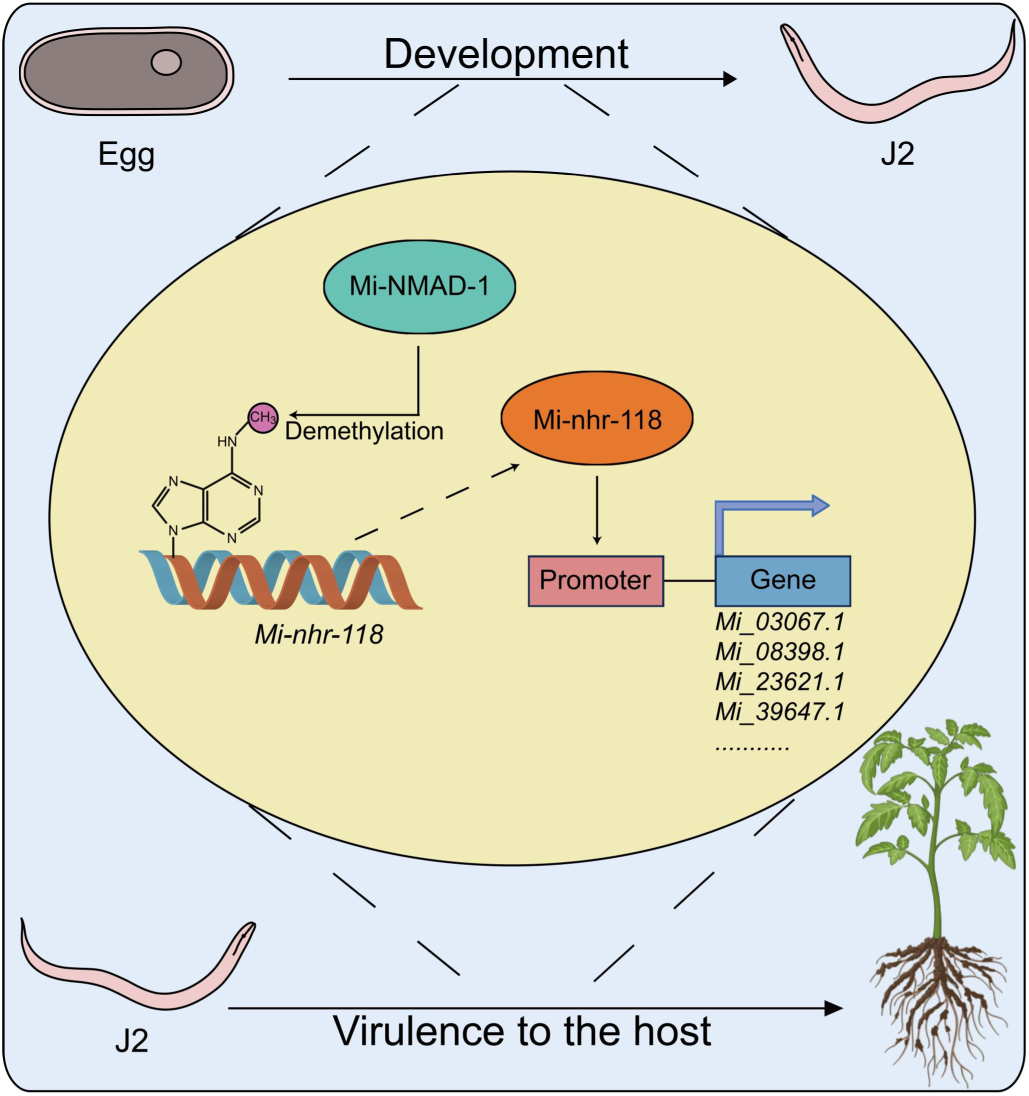
Model of Mi-NMAD-1–mediated regulation of nematode development and parasitism. The demethylase Mi-NMAD-1 regulates the transcription factor Mi-nhr-118, which in turn controls multiple downstream genes involved in development and parasitism, thereby modulating nematode growth and infectivity. The image of tomato was generated by BioRender.

## Materials and methods

### Nematode culture and embryo collection

The nematodes used in this study were purified strains previously maintained in our laboratory, including WHF4-1 (*M. incognita*), LongYF2-1 (*M. javanica*), and SYMaF1-2 (*M. arenaria*). Nematode-inoculated plants were grown in a greenhouse at 28 °C, 60% relative humidity, and a 16:8 h light–dark cycle for two months, after which nematodes at different developmental stages were harvested for experimental use. Nematodes at different developmental stages (egg, pre-J2, par-J2, par-J3/J4, and female) were collected as previously described ^23^. To collect RKNs embryonic cells, adult females were ruptured using fine-tipped forceps to release the reproductive organs. Approximately 500 µL of sterile water containing 0.01% Triton X-100 was added, and the suspension was pipetted several times to separate the cuticle from the embryonic cells. After natural sedimentation for about 10 seconds, the supernatant containing embryonic cells was collected. This procedure was repeated several times to maximize embryo yield.

### Sequence analysis, alignment and phylogenetic tree construction

The nucleotide and amino acid sequences of *Mi-nhr-118* were obtained from our previously published genome data ^23^. Protein domain annotation was performed on the Conserved Domain Database (CDD) ^37^. The orthologous proteins of *Mi-nhr-118* were identified from WormBase database using BLASTP tool with e-vaule < 10^−15^. Multiple sequence alignment of orthologous proteins was performed by mafft (v. 7.525) ^38^ and trimmed by trimal (v. 1.5.0) ^39^. The phylogenetic tree was constructed using iqtree (v. 2.3.6) ^40^ with the maximum likelihood (ML) method, incorporating 1000 bootstrap replicates, and visualized using iTOL ^41^.

### RNA extraction, cDNA synthesis, and cloning of *Mi-nhr-118*

Total RNA from nematodes at different developmental stages was extracted separately using TranZol Up (Cat. No. ET111; TransGen Biotech, Beijing, China) and quantified with a Qubit™ 4 Fluorometer (Invitrogen, Thermo Fisher Scientific, Waltham, MA, USA). Complementary DNA (cDNA) was synthesized from each RNA sample using the HiScript® RT SuperMix for qPCR (+gDNA wiper) (Cat. No. R323; Vazyme, Nanjing, China). The coding sequence of Mi-nhr-118 was amplified from cDNA using Ex Premier™ DNA Polymerase (Cat. No. RR370A; Takara, Dalian, China) and a specific primer pair, Mi-nhr-118_F and Mi-nhr-118_R (Supplementary Table 1).

### Determining the gene relative expression levels by RT-qPCR

The expression levels of target genes at different developmental stages were quantified using RT-qPCR. The *M. incognita* actin gene (Minc3s04507g36431, PRJEB8714) was used as an internal reference, with specific primers Mi-actin_F and Mi-actin_R (Supplementary Table S1). RT-qPCR was performed with Taq Pro Universal SYBR qPCR Master Mix (Cat. No. Q712; Vazyme, Nanjing, China) on an ABI QuantStudio™ 5 Real-Time PCR System (Thermo Fisher Scientific, Waltham, MA, USA). Relative gene expression levels were calculated using the 2^−ΔΔCt^ method.

### dsRNA synthesis and *in vitro* RNAi assays

Double-stranded RNA (dsRNA) fragments corresponding to the target genes and the green fluorescent protein (*gfp*) gene (used as a control) were synthesized *in vitro* using the T7 RNAi Transcription Kit (Cat. No. TR102-01; Vazyme, Nanjing, China) with the primers listed in Supplementary Table S1. The concentration of dsRNA was determined using a Qubit™ 4 Fluorometer (Invitrogen, Thermo Fisher Scientific, Waltham, MA, USA). To examine the effects of target gene silencing on embryonic development, RNA interference (RNAi) was performed on early embryos within female nematodes. Approximately 5,000 eggs were incubated with dsRNA (1 μg/μL) for 60 h at 25 °C on a rotator, followed by thorough washing with RNase-free water (Cat. No. P071-01; Vazyme, Nanjing, China). Subsequently, 100 eggs were carefully selected and transferred to a 96-well tissue culture plate (Cat. No. TCP001896; BIOFIL, Guangzhou, China), with 20 eggs per well (five wells in total). The plates were incubated in the dark at 25 °C and 70% relative humidity. Egg development was monitored and recorded every 24 h using an IX71 inverted microscope (Olympus, Tokyo, Japan). The hatching rate was calculated as the proportion of successfully hatched eggs relative to the total number of eggs. Treatment with dsGFP served as a negative control. The remaining treated eggs were used for transcriptome library construction. Each gene was biologically replicated three times.

### Generation of transgenic tobacco plants

Nuclear transgenic tobacco plants expressing hd-RNAi were generated as previously described ^42^. To construct a nuclear transformation vector for hairpin RNA (hd-RNAi) expression, target DNA fragments were first amplified using the primer pair Mi_gene-XhoI/Mi_gene-BglII and cloned as an XhoI/BglII fragment into the similarly digested vector pUC-RNAi (containing the StGA20 intron), yielding plasmid pUC-RNAi+Mi_gene-Up. Subsequently, target DNA fragments were amplified with the primer pair Mi_gene-EcoRI/Mi_gene-BamHI and inserted as an EcoRI/BamHI fragment into pUC-RNAi+Mi_gene-Up to generate plasmid pUC-RNAi+Mi_gene. Finally, the pUC-RNAi+Mi_gene plasmid was excised with XhoI and EcoRI and subcloned into the nuclear transformation vector pHW25, resulting in the final construct pHW25+Mi_gene (Supplementary Table S2).

Nuclear transgenic tobacco plants were generated using the Agrobacterium-mediated transformation method ^43^. Nicotiana tabacum cv. Petite Havana SR1 plants were cultured aseptically on Murashige and Skoog (MS) medium containing 20 g/L sucrose, 10 g/L agar, and 4.4 g/L MS salts with vitamins (Cat. No. PM1011-50L; Coolaber, Beijing, China). Young leaves from 4-week-old plants were cut into 5 × 5 mm pieces and infected with *Agrobacterium tumefaciens* strain EHA105 harboring plasmid pHW25+Mi_gene. After 48 h of co-cultivation in the dark at 28 °C, infected explants were transferred to RMOP medium (30 g/L sucrose, 6 g/L agar, 500 mg/L carbenicillin, 100 mg/L kanamycin, 1 mg/L 6-benzylaminopurine, and 0.1 mg/L 1-naphthaleneacetic acid) and incubated at 25 °C under a 16/8 h light–dark cycle (25 μmol·m⁻²·s⁻¹) for 2–3 weeks until shoot regeneration. Resistant shoots were excised and transferred to the rooting medium (MS medium supplemented with 0.1 mg/L 1-naphthaleneacetic acid and 100 mg/L kanamycin). Rooted plantlets were acclimated in soil after two weeks. Transgenic lines were verified by PCR detection of the XhoI/EcoRI fragment one month later, and seeds were collected from confirmed T₀ plants. T₁ seedlings, selected on MS medium containing kanamycin, were analyzed by qRT-PCR using gene-specific primers (Supplementary Table S2) to assess transcript abundance.

### Nematode infection assays

RNAi transgenic tobacco T₁ lines expressing dsRNA of target genes, along with control plants (WT and dsGFP), were used for nematode parasitism assays. One-month-old T₁ plants were grown in pots containing 200 g of a soil–sand mixture (3:1). Each line consisted of ten biological replicates. Plants were inoculated with approximately 1,000 freshly hatched pre-J2s of *M. incognita*, *M. arenaria*, *or M. javanica* and maintained in a growth chamber at 25 °C under a 16/8 h light–dark cycle (25 μmol·m⁻²·s⁻¹). Seventy days after inoculation, roots were gently washed to remove soil, and the number of galls, egg masses, and root fresh weight were recorded. Egg masses collected from different lines were used to assess hatching rates at 25 °C and 70% relative humidity. Statistical significance between treatments was determined using independent samples t-tests. Since we used 1,000 J2s for infection and collected data at 70 dpi, the control plants typically developed large, multi-generational root galls. These galls were too numerous and fused to be accurately counted, meaning the actual number was much higher than the recorded count.

### RNA-seq library construction and data analysis

Transcriptome library construction and sequencing were generated as previously described ^44^. Total RNA was extracted from the remaining dsRNA-treated eggs using the TransZol Up Plus RNA Kit (TransGen Biotech, Beijing, China). RNA integrity and concentration were evaluated with the RNA Nano 6000 Assay Kit on an Agilent 2100 Bioanalyzer (Agilent Technologies, Santa Clara, CA, USA). RNA-seq libraries were prepared from 2 µg of total RNA per sample using the VAHTS Universal V8 RNA-seq Library Prep Kit for Illumina (Cat. No. NR605-02; Vazyme, Nanjing, China). After quality assessment and cluster generation, sequencing was performed on an Illumina platform to produce approximately 6 Gb of 150 bp paired-end reads per library. Two biological replicates were included for each sample.

Clean reads were obtained using fastp (v0.23.2) ^45^ to remove adapter sequences and low-quality bases. The filtered reads were aligned to the *M. incognita* reference genome using HISAT2 (v2.2.0) ^46^ with a corresponding Gene Transfer Format (GTF) annotation file. Gene-level read counts were generated with HTSeq (v2.0.5) ^47^ and normalized using DESeq2 (v1.42.0) ^48^ to obtain expression values in normalized counts per million. Differentially expressed genes (DEGs) were identified using DESeq2, with significance thresholds of adjusted p < 0.05 and |log₂(fold change)| > 1. Functional annotation was performed by EggNOG (v2) ^49^ enrichment analysis of DEGs was performed based on Gene Ontology (GO) and Kyoto Encyclopedia of Genes and Genomes (KEGG) annotations. GO enrichment analysis was performed by using TBtools-II ^50^.

### Western blot

The Western blot procedure followed our previously published protocol ^5^ with minor modifications. Briefly, nematode eggs or J2s were treated with 1% sodium hypochlorite and purified by centrifugation through 35% sucrose. The nematodes were washed with PBS, flash-frozen in liquid nitrogen, and ground into a fine powder. After thawing, 50 µL of the supernatant was collected and mixed with 5 µL β-mercaptoethanol and 5 µL protein loading buffer, then boiled for 10 min and centrifuged at 12,000 rpm for 5 min. The resulting supernatant was used for SDS-PAGE. After electrophoresis and transfer onto a nitrocellulose membrane, the blot was incubated with the Mi-nhr-118 primary antibody (1:500) for 2 h at room temperature, followed by incubation with the secondary antibody and subsequent detection.

### ChIP-seq and analysis

ChIP-seq library construction and sequencing were performed as previously described. Approximately 50 µl of eggs or pre-J2s were collected from infected tomato roots, purified, and washed with pre-chilled PBS. Cross-linking was carried out by adding 38% formaldehyde and rotating at room temperature for 30 min, followed by quenching with 200 mM glycine. After centrifugation, the pellets were washed with cold 2× Nuclei Purification Buffer and homogenized using a rapid tissue grinder (65 Hz, 30 s × 5 cycles). Nuclear fractions were recovered by differential centrifugation and resuspended in ChIP lysis buffer. The lysates were washed sequentially with ChIP Wash Buffer and pre-chilled RSB buffer containing 0.1% Tween-20 to remove mitochondrial DNA. Nuclear pellets were split into two aliquots and subjected to chromatin fragmentation using a Covaris ultrasonicator (8 cycles, 30s on / 30s off). DNA was resuspended in RNase-free water, treated with micrococcal nuclease at 37°C for 30 min, and the reaction was terminated with EGTA. An aliquot was reserved as Input. The remaining chromatin was adjusted to FA buffer conditions, supplemented with RNase-free water, and incubated overnight at 4°C with the anti–Mi-nhr-118 antibody. Protein A/G magnetic beads (Cat# PB101-01, Vazyme, Nanjing, China) were used to pull down the antibody. ChIP-seq libraries were prepared from 5 µg of DNA per sample using the VAHTS Universal DNA Library Prep Kit for Illumina V3 (Cat# ND607-02, Vazyme). After library quality assessment and cluster generation, each library was sequenced on the Illumina platform to produce ∼6 Gb of 150 bp paired-end reads. Two biological replicates were generated for each sample.

Clean reads were aligned to the *M. incognita* genome ^23^ with Bowtie2 (v2.5.3) ^51^. Low-quality alignments (MAPQ < 10) were removed using Samtools (v1.19.2) ^52^, and PCR duplicates were filtered with Sambamba (v1.0.1) ^53^. Peak calling was performed with MACS2 (v2.2.9.1) ^54^, and peak-associated genes were annotated using ChIPseeker (v1.36.0) ^55^. De novo motif enrichment was conducted with HOMER (v4.11.1) ^56^ under default settings. Reproducible egg- and J2-specific binding peaks were defined as regions with >50% overlap between biological replicates.

## Data Availability

All sequencing data generated in this study have been deposited in the NCBI Sequence Read Archive (SRA). RNA-seq data related to demethylase HIGS are available under accession numbers SRR22190500–SRR22190505 and SRR22190516–SRR22190520. RNA-seq and ChIP-seq data for Mi-nhr-118 and its target genes have been deposited under BioProject PRJNA1362409, comprising 3 BioSamples and 46 SRA datasets.

## Notes

### Competing Interest Statement

The authors have declared no competing interest.

